# Prediction and annotation of alternative transcription starts and promoter shift in the chicken genome

**DOI:** 10.1101/2024.12.04.626748

**Authors:** Valentina A Grushina, Ivan S Yevshin, Oleg A Gusev, Fedor A Kolpakov, Sergey S Pintus

## Abstract

The promoter shift is a long known phenomenon which implies changes in the coordinates of a transcription start site (TSS). Analysis of CAGE experiments allows one to assess the impact and statistical significance of the shift. The promoter shift can be associated not only with the change of stages in early development or tissue differentiation, but also with the environmental influence on the cells. Differential promoter usage suggests non-constitutive expression activity of the regulated gene, which, in turn, suggests the usage of few sharp promoters. On the other hand, the housekeeping genes are assumed to demonstrate stable levels of expression while using multiple broad promoters. Additionally, the housekeeping genes are assumed to be normally expressed in most of the tissues. Our results show that many ubiquitously expressed genes use single sharp promoters and are subject to the significant promoter shift, which implies an additional level of complexity to the problem of definition of a housekeeping gene.

The reaction of genes in cells and tissues to external stimuli is often studied in the context of differential gene expression. Our study employs a different approach of differential promoter usage, which takes the genes whose promoters undergo significant promoter shifting as the signatures of tissue response and phenotypic effect. Our results suggest that the difference in growth rate of chickens is controlled by the rate of metabolism of nutrients via the differential usage of promoters of the relevant genes.

## Introduction

The analysis of promoter regions of genes often reveals differences from the existing annotation, which can be caused both by the insufficient accuracy of the existing annotation and the existence of transcripts previously unknown and activated under certain non-standard conditions [Link et al, 2018; Warren et al, 2020]. Taking into account the data from CAGE experiments would make it possible to better identify the initial positions of such transcripts, which allows us to determine the position of the start of transcription, as well as to identify the events of the promoter shift [Haberle et al, 2014]. Such insights would provide a better understanding of the mechanisms of transcriptional regulation in the chicken genome and improve the quality of Gallus gallus genome annotation.

Conventional pipeline for analysis of CAGE sequencing data involves aggregation of the start positions of the mapped reads into 0 bases wide intervals starting from a certain coordinate on a chromosome, known as CAGE tag start sites (CTSS), each representing a number of cases when transcription started exactly from that genome coordinate. However, gene transcription does not usually start from the very single base – one would rather observe a variety of CTSS, forming a peak or a cluster within a short genomic interval. This interval is referred to as the transcription start site (TSS) and all the CTSS within it are usually distributed around the most expressed CTSS which is often referred to as the TSS peak center or the dominating CTSS [Forrest et al, 2014].

When comparing samples from two different tissues or experiment conditions, one may observe that the distribution of the CTSS tags along the TSS is not unimodular, but rather demonstrates a mixture of two different distributions each being characteristic to the specific tissue or the certain experiment condition. Such a shift in transcription start was supposed to take place due to differential promoter usage and was subsequently termed the ‘promoter shift’ or ‘TSS shifting’ [Haberle et al, 2014].

The phenomenon of the promoter shift itself was discovered much earlier as some mutations in the transcription factor TFIIB enabled it to shift the transcription start site [Pinto et al, 1994; Faitar et al, 2001]. In CAGE experiments, the promoter shift was originally observed in differentiating cells of a zebrafish embryo in the course of the maternal to zygotic transition [Haberle et al, 2014]. Recently, altered promoter usage in the same tissue in adults was associated not only with differential gene expression [Gacita et al, 2020], but also with response to an environmental change [Rosikiewicz et al, 2021].

In this study, we used sequencing data from CAGE experiments on various chicken tissues, including muscles, kidneys, liver, brain, and others. This provided us with the opportunity to annotate enhancers and alternative initial transcription positions that were active in various tissues. This approach ensures the completeness of the annotation, taking into account the differences in the gene expression in different tissues.

A comprehensive evaluation of the biological implications of promoter shift necessitates exhaustive annotation of downstream genes to ensure accurate interpretation of their functional relevance. Nowadays, RefSeq and Ensembl are two major gene annotation systems which are mostly used. Also available is the gene annotation used in the UCSC genome browser which is currently

Data from CAGE experiments are traditionally used to determine the start coordinates of gene models. Nevertheless, the TSS peaks imputed from CAGE data are not always consistent with the traditional annotation systems such as RefSeq or Ensembl. The discrepancies might also arise due to variability of completeness of gene annotation among genome sequencing projects. Particularly, the chicken genome annotation seems way less complete in comparison to that of the human genome. Thus, some part of the predicted TSS might relate to existing genes, not currently included in the genome annotation. On the contrary, some annotated genes whose expression is below detection might not have any TSS predicted proximate to their start coordinates.

In this study, we predicted transcription start sites (TSS) in the chicken genome and evaluated their expression based on data obtained from CAGE experiments on growing chickens and estimated the regulatory potential of the detected promoter regions. We also evaluated the significance of promoter shift as between tissues, as well as between slow growing and fast growing contrast groups for each of the tissues studied.

## Methods and Algorithms

### Public CAGE data

For this study we used CAGE data on samples from 24 chickens of F2 generation from a cross between the Russian White and Cornish breeds. The birds were equally divided into the two categories based on the average daily weight gain – slow growing (12 chickens) and fast growing (12 chickens). From each bird leg, breast, brain, liver, heart, and kidney tissues were sampled.

The CAGE-seq paired end reads (DNBSEQ-G400) from 6 tissues, taken from 24 chickens, (12 slow growing and 12 fast growing) at the age of 9 weeks: heart, kidney, leg, liver, brain, breast were downloaded from Chicken GeneTech database (URL https://chicken.biouml.org/downloads/ChickenResearch2023/CageSeq/raw_data/).

For validation of our results we obtained CAGE reads of chicken tissue samples from the FANTOM5 project [Kawaji et al, 2017] website, remapped them to the galGal6 reference genome, aggregate CTSSs and clustered TSS peaks using the same approach as for the F2 breed samples.

### Reference genome

We used the the GRCg6a (galGal6) RefSeq reference genome assembly as available from NCBI (ID GCF_000002315.6)

### Reference annotations

The Ensemble and NCBI RefSeq genome annotations available for the corresponding genome assemblies, namely Ensembl release 106 and NCBI RefSeq Gallus gallus annotation release 104 (GCF_000002315.5), were used as a reference for annotating, comparing and validating TSS and CTSS coordinates.

### Read mapping

CAGE paired end reads were aligned against the reference genomes using STAR RNA-seq aligner [Dobin et al, 2013] (version 2.7.11b), separately for each sample.

### CTSS aggregation and annotation

The start positions of the mapped CAGE reads were aggregated into CTSS similarly to the approach used in the FANTOM5 project, which employed filtering with SAMtools, conversion and subsequent aggregation with BEDtools [Lizio et al, 2017a]. The resulting CTSS were converted from BED format to the native CAGEr format (chromosome, tag start, strand, number of reads supporting the tag). The CTSS were then loaded in the CAGEr package [Haberle et al, 2014], which was used for their subsequent annotation with the RefSeq gene models.

### Power law based normalization of the CTSS expression values

For the normalization of the CTSS we used the method based on power law distribution of CTTS values [Balwierz et al, 2009] as implemented in the CAGEr package. We used two sets of normalization parameters – robust and permissive. For the robust normalization we used -1.2 as the slope of the power law distribution of the CTSS values in log-log coordinates and 5e +04 as the referent number of the CTSSs. For the permissive normalization we relaxed the referent number of CTSSs to 1.2e +07 while leaving the slope parameter the same as for the robust case.

### Contrasts and TSS peak clustering

We clustered the CAGE tags into the CTSS in a two-step manner, as the standard CAGEr pipeline suggests. We divided all samples into biologically relevant groups, generated group-specific TSS peaks and then aggregated those peaks into the consensus clusters. Such hierarchical grouping allowed us to estimate the promoter shift between different contrasts. For the set of contrasts we divided all samples into six tissue-specific groups, each containing 24 samples of a certain tissue type. Then we subdivided each set of tissue samples into two groups of slow growth and fast growth

For clustering on the group level, CAGEr’s implementation of the DISTCLU algorithm was used. For DISTCLU clustering all CTSSs below the threshold of 1 transcript per kilobase million reads (TPM < 1) were filtered out. Additionally, the singleton CTSS having less then 5 reads starting from them were removed. The maximal distance between two neighboring CTSSs in a cluster was 20 nt. The resulting TSS peaks were also annotated with the RefGene set and the CAGE tags were mapped against them.

To identify the events of significant promoter shift we applied the CAGEr’s Kolmogorov-Smirnov test of the differential CAGE tag usage to the unnormalized CTSS counts in the consensus clusters.

To identify the promoter shift events in ubiquitously expressed genes as between tissues as well as between slow and growing chickens in the tissues under study we used robust normalization. To identify the promoter shift events between genes which included ones that were specific to certain tissues we used the permissive normalization (see above).

### Gene set enrichment analysis

We performed gene set enrichment analysis using the exact Fisher test with FDR correction of P values PANTHER service against the GO Slim database of biological processes [Thomas et al, 2022; Mi et al, 2019] and the Bioconductor package clusterProfiler [Xu et al, 2024] against the KEGG Pathway database [Kanehisa et al, 2023].

## Results

### Tissue characteristic promoter regions

The clustering of CAGE tags in tissue groups resulted in six roughly equal sets of tag clusters mostly annotated to exonic and intergenic regions (ST1). Out of all genes which annotated the tag clusters (totally 1215 genes), most were detected in all tissues (1187 genes) and tissue specific genes, which TSSs were detected only in one tissue, were pretty few (16 genes) (Table 1). Moreover, nearly 90% of the genes (1091 out of 1215) had only one active TSS in all six tissues and only a few had more than 3 (Table 2).

**Table 1.** Tissue specific genes of the tissue characteristic tag clusters.

| Tissue | Tissue specific genes |
| --- | --- |
| brain | AVP, SLC38A2, VAMP1 |
| breast | KLHL31, MYH1E |
| kidney | CYP2AC1, ETNK1, FCGBPL |
| heart | C7, LPL, RRAD, RYR2, TPM4 |
| leg | CCLI5, ZNF106 |
| liver | PHGDH |

**Table 2.** Number of annotated tissue characteristic TSSs per gene and number of genes having that many TSSs.

| Number of TSSs in a gene | Number of genes |
| --- | --- |
| 1 | 1091 |
| 1 – 2 | 2 |
| 2 | 75 |
| 3 | 11 |
| 3 – 4 | 1 |
| 4 | 3 |
| 8 | 1 |
| 26 | 1 |
| 34 | 1 |

On the level of the tag clusters, most of the TSSs were also active in all six tissues, namely 2350 out of 2387 non-overlapping genomic intervals. Notably, the intergenic TSSs followed the similar statistic – 1027 out of 1035 non-overlapping intergenic tag clusters were as well active in all six tissues.

The ubiquity of the active TSSs among all studied tissue samples suggested that the gene expression differentiation between tissues did not happen in a ‘switch-on/switch-off’ manner, but was rather based on a variety of observable levels of mRNA.

### Ubiquitously active promoter regions

The aggregation of consensus clusters from tissue characteristic TSSs allowed us to focus on the set of the promoter regions which were active in all tissues. The set of the TSSs narrowed to 507 genomic intervals, out of which 308 were annotated in galGal6 RefSeq and nearly each of them (306) was the single TSS region of its annotated gene (ST2). This prevalence of the single-promoter genes allowed us to validate the consensus clusters as the TSSs of the putatively regulated genes, according to the model of a core promoter described by Kadonaga [Kadonaga, 2011]. The model suggested the narrow TSSs of the single-promoter genes and wide TSSs of genes with multiple TSSs. Indeed, the median interquartile range of the consensus TSSs, including all the intergenic and both intronic, was only 6 nt (Figure 1). The exceptional TSSs were annotated to the LOC112533555 gene (an ortholog of the olfactory receptor gene OR14J1), and the LOC112533599 gene (RNA28S18, 28S ribosomal RNA 18). Both genes had 5 comparatively wide (35 – 1004 nt) TSSs each, being consistent with the Kadonaga’s notion on wide multiple promoters and sharp single promoters.

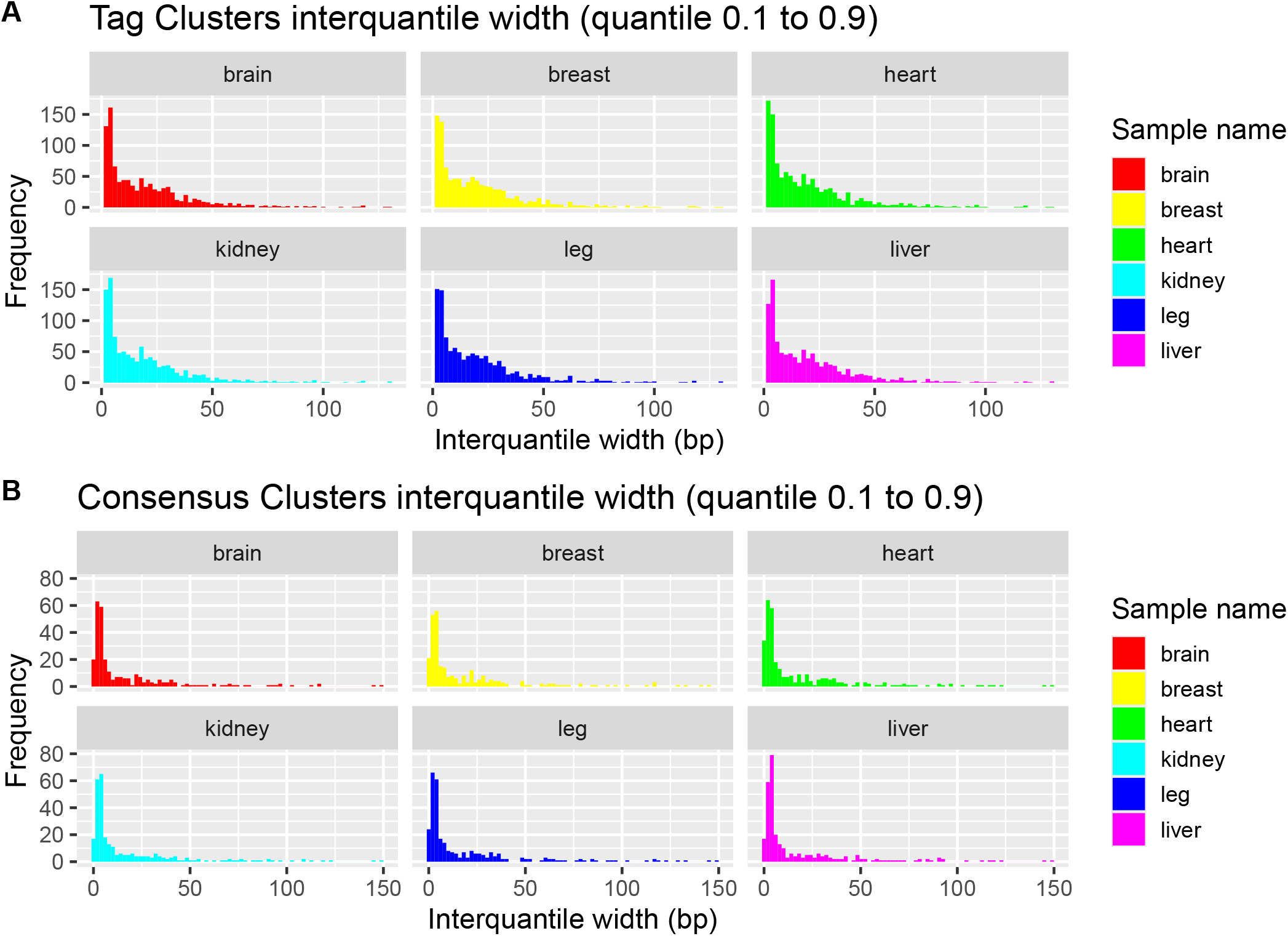

### Housekeeping status of the ubiquitous and tissue characteristic genes

Notably, the set of genes which annotated the ubiquitously active promoters rather weakly intersected with the set of the prominent chicken housekeeping genes identified in [Lizio et al, 2017b] – out of 308 ubiquitously expressed genes only 68 (22%) intersected with the 1256 gene symbols of the housekeeping set.

Similarly, the vast majority (92%) of the tissue characteristic TSSs were the single promoters of their genes. The rest 8% were expressed in all tissues and comprised of 90 genes with 2 or 3 promoters and only 3 genes, namely LOC112532777 (carbamoyl-phosphate synthase 1, CPS1), LOC112533555 (olfactory receptor 14C36-like), and LOC112533599 (28S ribosomal RNA 18, RNA28S18), had 8, 26 and 34 promoters each. Expectantly, only 282 out 1091 tag characteristic genes (26%) expressed in all 6 tissues intersected with the aforementioned housekeeping genes set of [Lizio et al, 2017b].

The chicken gene set of Lizio et al followed the criterion of ubiquitous expression, low tissue variance, and no exceptional expression [Eizenberg & Levanon, 2013]. From one hand, our gene set followed the principle of ubiquity, but on the other hand, Kadonaga’s model of housekeeping genes suggested multiple broad promoter regions per gene [Kadonaga, 2011], while we identified genes which mostly had sharp and single promoters.

### Shifting promoters between tissues

Although the promoter region activity did not exhibit the tissue specificity in terms of presence or absence, we found a significant shift in the usage of the TSS regions between different tissues. Interestingly, the usage of more than half of the consensus TSSs were significantly different between any pair of tissues (ST2). Expectantly, the leg and breast tissues were the most proximate to each other as the number of the TSSs affected by significant promoter shift between them was the least (Figure 2, Table 3).

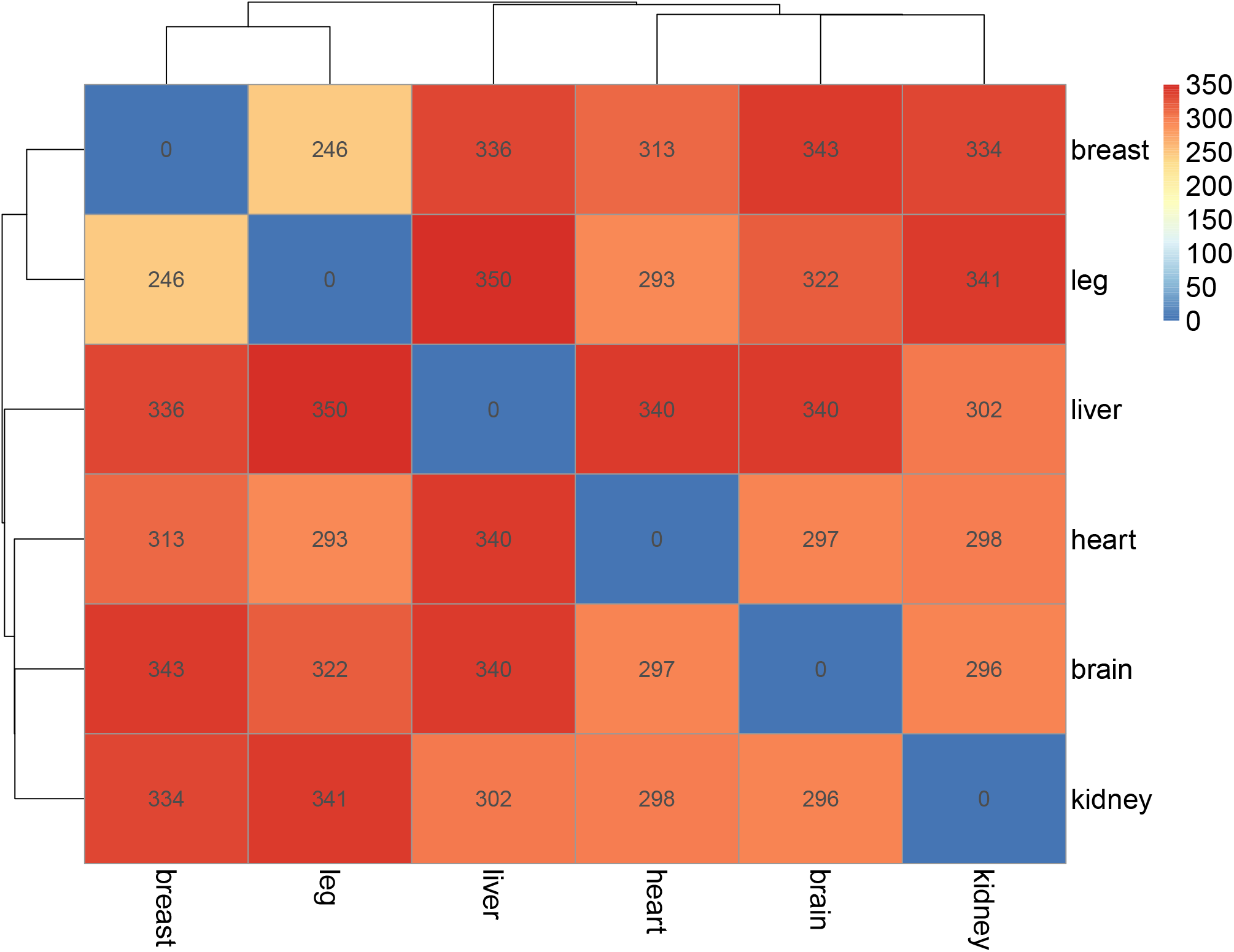

**Table 3.** Numbers of TSSs exhibiting significant promoter shift (FDR < 0.01) between tissue samples in each pair. Each summary number in the column ∑ is the total distance of the tissue from all other tissues.

| | brain | breast | heart | kidney | leg | liver | $\Sigma$ |
| --- | --- | --- | --- | --- | --- | --- | --- |
| brain | 0 | 343 | 297 | 296 | 322 | 340 | 1598 |
| breast | 343 | 0 | 313 | 334 | 246 | 336 | 1572 |
| heart | 297 | 313 | 0 | 298 | 293 | 340 | 1541 |
| kidney | 296 | 334 | 298 | 0 | 341 | 302 | 1571 |
| leg | 322 | 246 | 293 | 341 | 0 | 350 | 1552 |
| **liver** | 340 | 336 | 340 | 302 | 350 | 0 | 1668 |

### Shifting promoters between the same tissues in slow and fast growing chickens

Interestingly, most of the genes, which TSSs exhibited significant promoter shifts between the same tissues in slow and fast growing chickens were the same in all 6 tissues (ST3). These intervals belonged to 32 genes which demonstrated significant enrichment of the signatures of protein cascade, negative blood coagulation, inosine monophosphate (IMP) metabolism and mitochondrial electron transport (ST4). Most TSSs in fast growing chickens were shifted not more than to 100 bp upstream or downstream of the TSS position in slow growing chickens. Nevertheless, few TSSs demonstrated promoter shift up to the distance of 300 – 400 bp. (Figure 3)

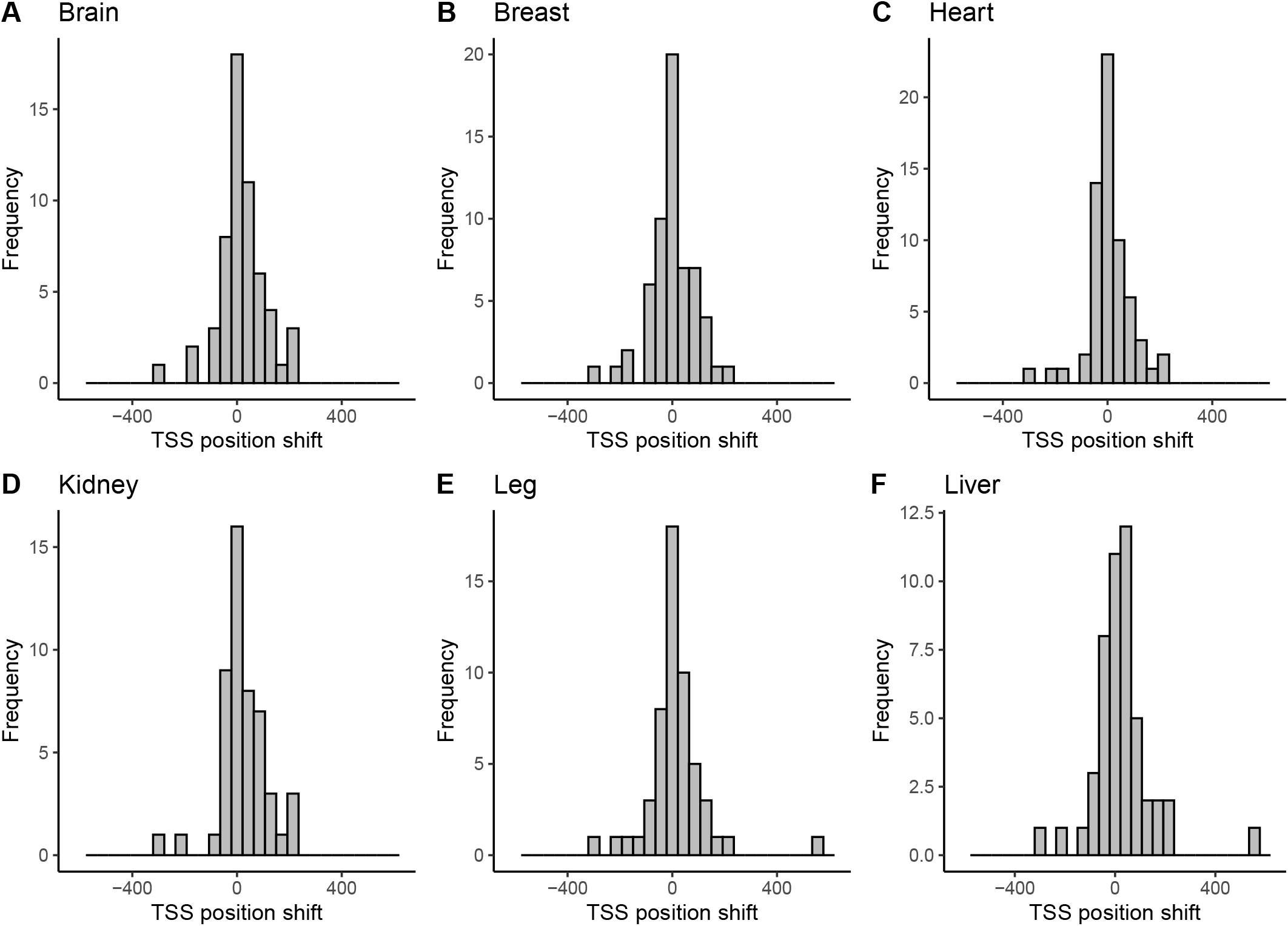

### Tissue specific promoter activity and shifting promoters

In order to assess the tissue specificity of promoter activity in slow and fast growing chickens we relaxed the normalization to 10^8^ total tags (see Methods and Algorithms) and recalculated the consensus clusters and promoter shifting events (ST5). The resulting permissive set of consensus clusters allowed us to identify tissue specific TSSs and their corresponding genes which underwent significant promoter shift between slow and fast growing chickens. Expectantly, on the tissue specific level, the promoter shift affected different pathways in different tissues. (Figure 4, ST6). Thus, the genes with promoter shifting were significantly enriched with signatures of fatty acid metabolism, lysosome and endocytosis only in brain, signatures of peroxisome and degradation of valine, leucine and isoleucine, as well as fatty acids, only in kidney, autophagy signatures only in breast and signatures of metabolism of glutathione, glycine, serine, threonine, glyoxylate and carboxylate only in liver.

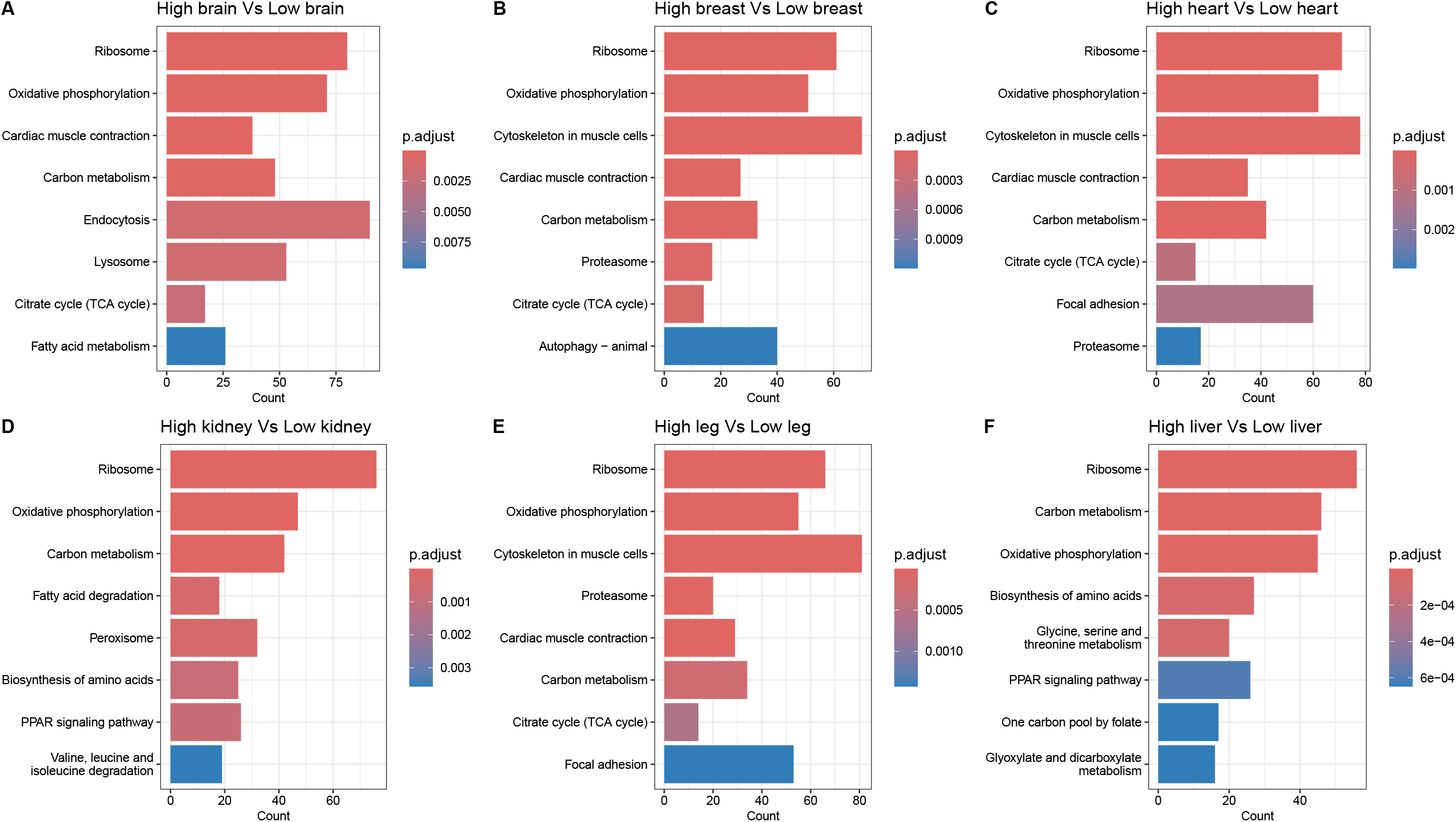

The patterns of presence and absence of the enriched KEGG pathways among genes with shifted promoters in different tissues formed a pattern that separated the liver and kidney tissues from the rest of the tissues under study (Figure 5).

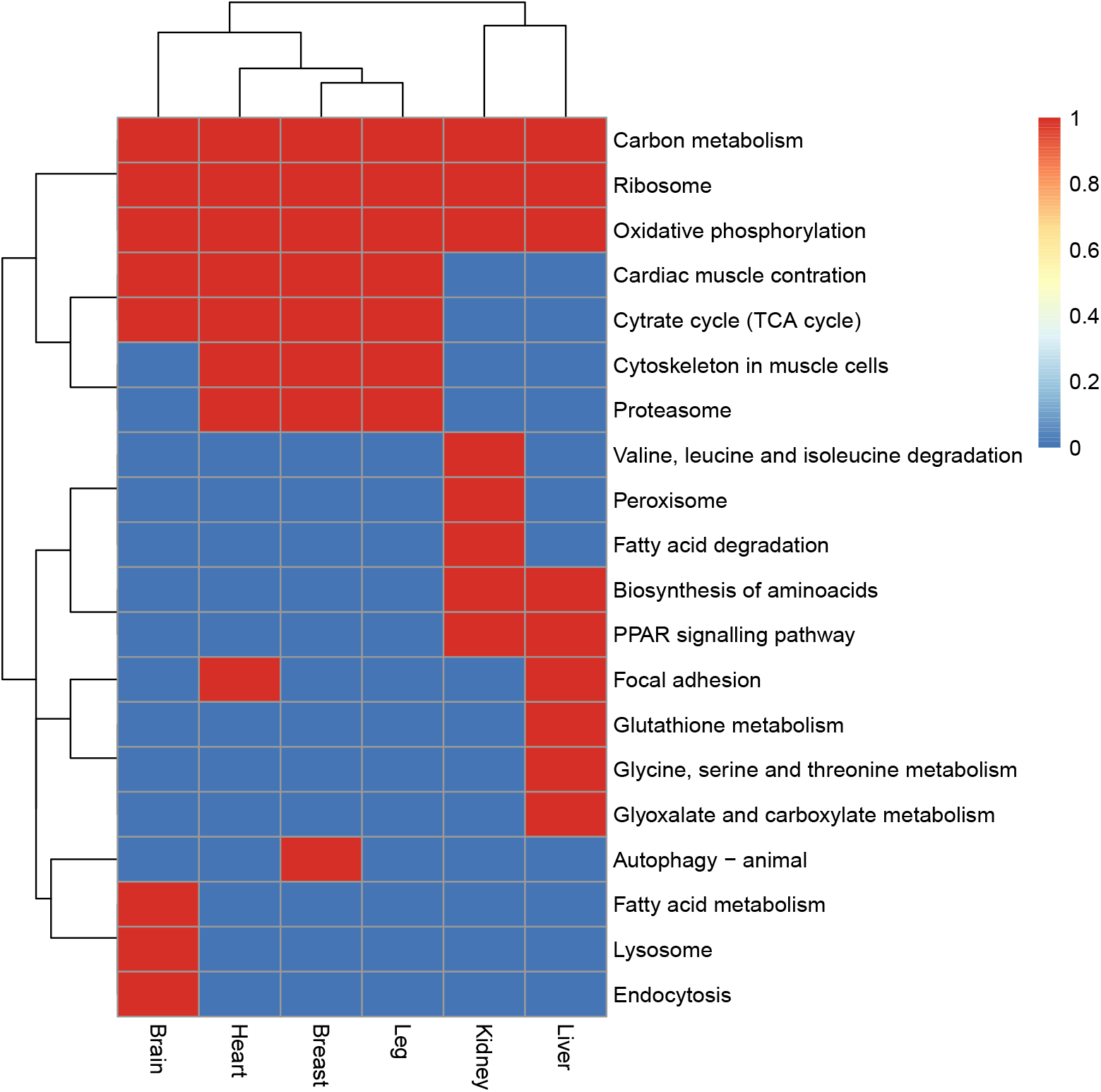

Expectantly, involvement of most of the enriched pathways in growth is associated with nutrition uptake and muscle tissue growth. Thus, lysosomal acid lipases are known to free fatty acids from lipids which enter the lysosome in the brain, particularly through endocytosis [Triverdi et al, 2020]. Interestingly, the autophagy signatures were significantly enriched only in the breast but not in the leg, given that both body sites are rich in skeletal muscle tissue. The glyoxylate cycle – a variation of citric acid cycle – has traditionally been associated with bacteria, protists, plants and fungi, but was also identified in chicken liver [Davis et al, 1990]. Peroxisomes of all species are known to participate in β-oxidation of fatty acids [Wanders & Waterham, 2006]. In mice, high-fat diets induced proliferation of peroxisomes [De Craemer at al, 1994] and chicken kidney were shown to express peroxisome proliferator-activated receptor (PPAR) genes [Meng et al, 2005]. Signatures of PPAR signaling itself were enriched not only in the kidney, but also in the liver. Curiously, the differentially regulated signatures of growth were enriched only in the kidney. Although lipogenesis in chickens have been shown to take place mainly in the liver [Cartoni Mancinelli et al, 2022] the overweight chickens are known to suffer from fatty liver and kidney syndrome which is associated with accumulation of fatty acids not only in liver but also in kidney [Whitehead, 1975]. It may be supposed that the fast growing chickens benefited from improvement of fatty acid degradation exactly in the kidney.

## Discussion

### Promoters width and count, their expression abundance and housekeeping status of the downstream genes

The core promoter gene model of Kadonaga [Kadonaga, 2011] makes a difference between regulated genes with few sharp promoters and housekeeping genes with multiple broad promoters. At the same time the concept of housekeeping genes proposed by Eisenberg and Levanon [Eisenberg & Levanon, 2013] suggests that housekeeping genes are ubiquitous. In present work we observed that the genes which were ubiquitous in tissue representation mostly had one sharp promoter. Additionally, we observed weak, although significant, intersections with the previous findings on housekeeping genes in chicken [Lizio et al, 2017b]. Therefore one might hypothesize that not all housekeeping genes are resilient to regulation and not all ubiquitously expressed genes can actually be termed housekeeping.

### Tissue specific gene regulation

Tissue specificity of gene expression is generally considered a quantitative, not qualitative metrics, i.e. some level of expression can be detected in some tissue even for a non-specific gene [Kryuchkova-Mostacci & Robinson-Rechavi, 2017]. Our results on promoter shift between tissues in the consensus set of the TSSs demonstrated that gene expression between tissues is also subject to tissue specific differential regulation of ubiquitously expressed genes. Particular tissue specific promoters are well known not only in animals [Meister et al, 2010] but also in plants [Zierer et al, 2022]. In our study we demonstrated that the tissue specific promoter shift is abundant among genes and tissues.

### Differential regulation of growth related genes and pathways

In our research we found pathways that supposedly were affected by differential regulation via the shifting of the promoters of the involved genes. Some of those pathways are consistent with the results of previous differential gene expression studies. Particularly, the involvement of the focal adhesion pathway was demonstrated for chicken legs in transition between stages of slow and fast growth [Xue et al, 2017]. Curiously, the growth hormone receptor pathway whose activation was previously associated with myoblast differentiation and muscle growth in chicken [Xu et al, 2019] was not the enriched term among the differentially regulated genes in our study suggesting that the differences between the slow growing and fast growing chickens were not the result of the myoblast differentiation on early development stage.

## Conclusion

In addition to the major transcription starts annotated in Refseq and Ensembl reference annotations, CAGE experiments reveal additional TSS. Our results demonstrate that a significant fraction of genes are affected by the phenomenon of transcription start shifts that manifest under specific conditions.

Our results suggest that the significant amount of genes which were abundant among the most of the tissues were subject to transcription regulation according to the strength and narrowness of their TSSs in contrast to the notion that ubiquitously expressed genes were mostly housekeeping and consequently possess broad promoters which made transcription of those genes resilient to regulation.

According to our results, the promoters of the aforementioned ubiquitously expressed genes were subject to promoter shift between tissues, suggesting different pathways of regulation of those genes in different tissues.

Besides the fine-tuning of tissue diversity, the effect of promoter shift participated in differential regulation of growth related genes in the same tissue but in contrast groups of growth rate. Our resulting signatures of differential regulation were enriched with pathway terms related to enhanced metabolism of nutrients, but not hormonal regulation of growth related promoters.

## Supporting information

Supplementary Table 1 - Tag Clusters by Tissue - Robust Set

Supplementary Table 2 - Consensus Clusters (Tissue Comparison) - Robust Set

Supplementary Table 3 - Consensus Clusters (Slow vs Fast Growing Comparison) - Permissive Set

Supplementary Table 4 - PANTHER GO Slim Enrichment Slow vs Fast Growing - Robust Set

Supplementary Table 5 - Promoter Shifting Between Slow and Fast Growing - Permissive Set

Supplementary Table 6 - KEGG Pathways Enrichment in Promoter Shifted Genes (Slow vs Fast Growing - Permissive Set)

